# Lung evolution in vertebrates and the water-to-land transition

**DOI:** 10.1101/2022.02.14.480340

**Authors:** Camila Cupello, Tatsuya Hirasawa, Norifumi Tatsumi, Yoshitaka Yabumoto, Pierre Gueriau, Sumio Isogai, Ryoko Matsumoto, Toshiro Saruwatari, Andrew King, Masato Hoshino, Kentaro Uesugi, Masataka Okabe, Paulo M. Brito

## Abstract

A crucial evolutionary change in vertebrate history was the Palaeozoic (Devonian ~400 million years ago) water-to-land transition, allowed by key morphological and physiological modifications including the acquisition of lungs. Nonetheless, the origin and early evolution of vertebrate lungs remain highly controversial, particularly whether the ancestral state was paired or unpaired. Due to the rarity of fossil soft tissue preservation, lung evolution can only be traced based on the extant phylogenetic bracket. Here we investigate, for the first time, lung morphology in extensive developmental series of key living lunged osteichthyans using synchrotron X-ray microtomography and histology. Our results shed light on the primitive state of vertebrate lungs as unpaired, evolving to be truly paired in the lineage towards the tetrapods. The water-to-land transition confronted profound physiological challenges and paired lungs were decisive for increasing the surface area and the pulmonary compliance and volume, especially during the air-breathing on land.

## Introduction

Lungs, the most important organ of the pulmonary complex, are rarely preserved in fossils, hindering direct evidence of how the earliest air-breathing vertebrates breathed air. So far, the evolutionary origin of the vertebrate lung has been narrowed down to the basal osteichthyans (Goujet, 2011; Tatsumi et al., 2016). However, since the knowledge about morphological and genetic development of the lung has been highly biased in amniotes, the original form of this evolutionary novelty has remained elusive. One hypothesis, formed and supported by studies on tetrapods (particularly mammals and birds), assumes that the lung evolved through a modification of the pharyngeal pouch (Kastschenko, 1887), as the lung bud develops at the pharyngo-oesophageal junction during embryonic development. Consequently, this view (Kastschenko, 1887; Kuratani and Tanaka, 1990) predicts that the primitive lungs appeared as bilaterally paired organs at the caudolateral part of the pharynx. Indeed, in embryology, lungs of living tetrapods have been mostly described as paired derivates of the respiratory tube, arisen from paired and small hollow swellings (Marshall Flint, 1990). Previous studies on amphibians have also proposed that the lung bud develop from paired rudiments of the ventral portions of the eighth pharyngeal pouches (Goodrich, 1931; Marcus, 1937; Perry et al., 2001). Additionally, the plesiomorphic state of lungs has been mostly described as paired organs (Funk, Lencer and McCune, 2020). On the other hand, another hypothesis does not constrain the evolutionary origin of the lung to the serial homologue of the pharyngeal pouch (Greil, 1913; Neumayer, 1930; Wassnetzov, 1932). In this view, although the possibility that the primitive lung developed on the pharyngeal endoderm is not excluded, the primitive lung is considered to appear on the floor of the pharynx, or more generally, on the floor of the foregut. This scenario does not predict bilaterally paired forms of primitive lungs.

Curiously, some living vertebrates display an unpaired organ (Cupello et al., 2015; Cupello et al., 2017a; Cupello et al., 2017b; Cupello, Clément and Brito, 2019; Lambertz and Perry, 2015; Lambertz et al., 2015), leaving the ancestral condition equivocal. The sister group to all other extant actinopterygians, the obligate air-breathing polypterids (Icardo et al., 2017), breath air using lungs, which have previously been described as a paired organ (Icardo et al., 2017; Geoffrey Saint Hilaire, 1802; Graham, 1997). However, in adult specimens of *Polypterus senegalus* the glottis only opens to the right sac and the left sac is connected to the right sac by a separate opening (Graham, 1997), raising old questions about its true paired condition. Among sarcopterygians, the unpaired lung of coelacanths is unequivocal. The living coelacanth *Latimeria chalumnae,* that inhabits moderate deep-water and makes gas exchanges only through gills, have an unpaired lung with no outline of a second bud at different developmental stages (Cupello et al., 2015; Cupello et al., 2017a; Cupello, Clément and Brito, 2019). In the extant sister group of all tetrapods (Amemiya et al., 2013), namely lungfishes, the three extant genera have lungs capable to uptake oxygen from the air. However, in the most basal one, the facultative air-breather *Neoceratodus forsteri,* the lung is described as unpaired (Greil, 1913; Graham, 1997; Grigg, 1965). In contrast with both South American and African lungfishes, the Lepidosirenoidea, that are obligated air-breathers and have a lung described as a ventral paired organ (16) like tetrapod lungs.

To follow lung evolutionary history in vertebrates, we analyzed primitive sequence of morphogenesis of lungs of key living osteichthyans (Figs. 1–6). Embryos, larvae, juveniles and adults of *P. senegalus, N. forsteri, Lepidosiren paradoxa* were examined. To compare the lung anatomy of osteichthyan fishes with tetrapods, we studied also an extensive developmental series of the living salamandrid *Salamandra salamandra,* from early and late larvae to juveniles before and after metamorphosis (Fig. 5). As salamanders are often considered to have retained plesiomorphic characteristics of tetrapod stance and locomotion (Pierce et al., 2020), we used them here also as a model for understanding lung evolution in tetrapods. Specimens of mentioned taxa were examined through x-ray microtomography, the unique effective non-invasive methodology to study their morphology and histology at a three-dimensional (sub) microscale. When possible, we proceeded also with dissections and the study of histological sections. We compare our results with the available information from the lung of fossil taxa, the coelacanths and salamanders (Cupello, Clément and Brito, 2019; Brito et al., 2010; Tissier, Rage and Laurin, 2017).

**Fig. 1.**
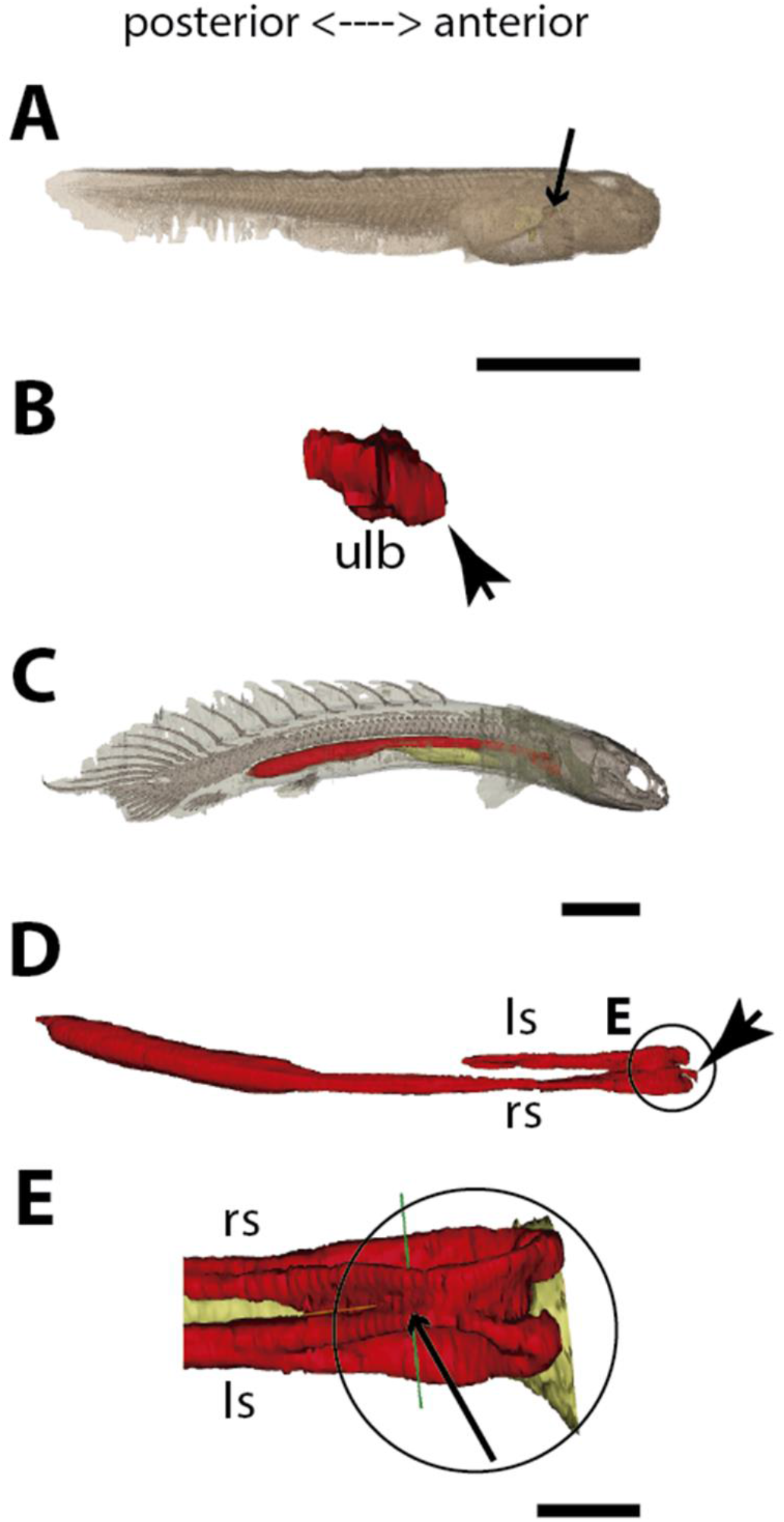
Three-dimensional reconstructions of the pulmonary complex of *Polypterus*. **(A)** Early embryo (9.3 mm) TL in right lateral view, **(B)** Isolated lung of the early embryo in dorsal view, **(C)** Juvenile (45 mm TL) in right lateral view, **(D)** Isolated lung of the juvenile in dorsal view, **(E)** Close-up of (D) highlighting the lung in ventral view and pointing out the region of the independent and secondary connection of the left sac to the right one by a lateral opening. Yellow, foregut including the stomach; red, lung. Black arrow in **(A)** pointing to the lung. Arrowheads in **(B)** pointing to the lung connection to the foregut and in **(D)** pointing to the pneumatic duct connection to the foregut. Black arrow in **(E)** pointing to the independent connection. Ls, left sac; rs, right sac; ulb, unpaired lung bud. Scale bars, 5.0 mm **(A)**; 0.075 mm **(B)**; 5.0 mm **(C, D)**; 1.0 mm **(E)**.

**Fig. 2.**
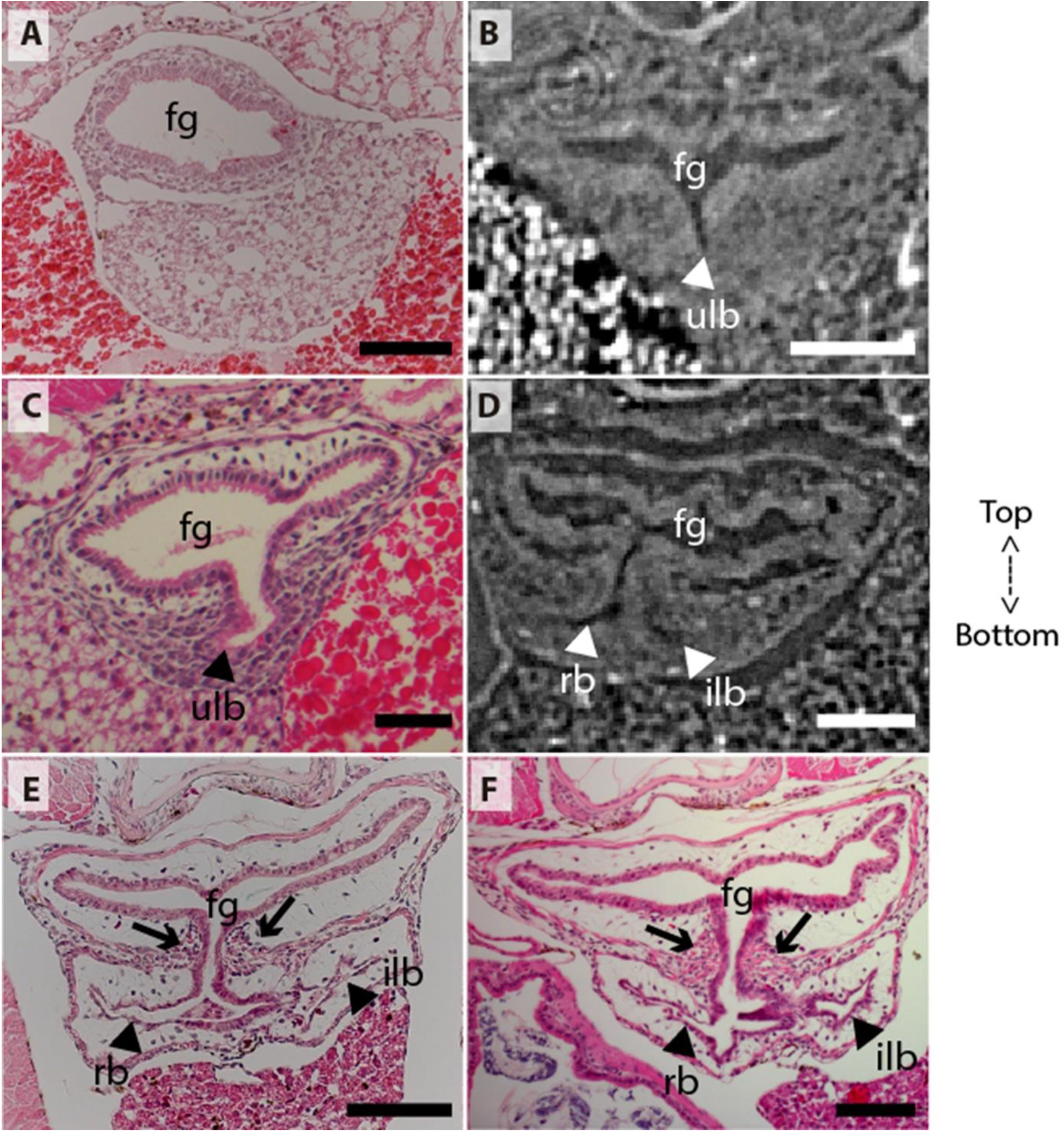
Coronal sections of the unpaired lung in the living actinopterygian fish *Polypterus senegalus*. **(A)** No lung bud in 8.0 mm TL specimen, **(B)** Origin of an unpaired lung bud in 8.5 mm TL specimen, **(C)** Unpaired lung bud in 9.1 mm TL specimen, **(D)** First register of an independent and lateral second lung bud in 12 mm TL specimen, **(E, F)** Independent and lateral second lung bud arising from the principal tube in 15.5 mm TL and 18 mm TL specimens. **(A, C, E–F)** Histological thin-sections. **(B, D)** Sections of synchrotron X-ray microtomography of the early embryo. Black and white head arrows pointing to the lumen of the unpaired lung buds; arrows pointing to the undifferentiated cells surrounding the glottis. Fg, foregut; ilb, independent lateral bud; rb, right bud; ulb, unpaired lung bud. Scale bars, 0.2 mm **(A, E)**; 0.1 mm **(B, D, F)**; 0.05 mm **(C)**.

**Fig. 3.**
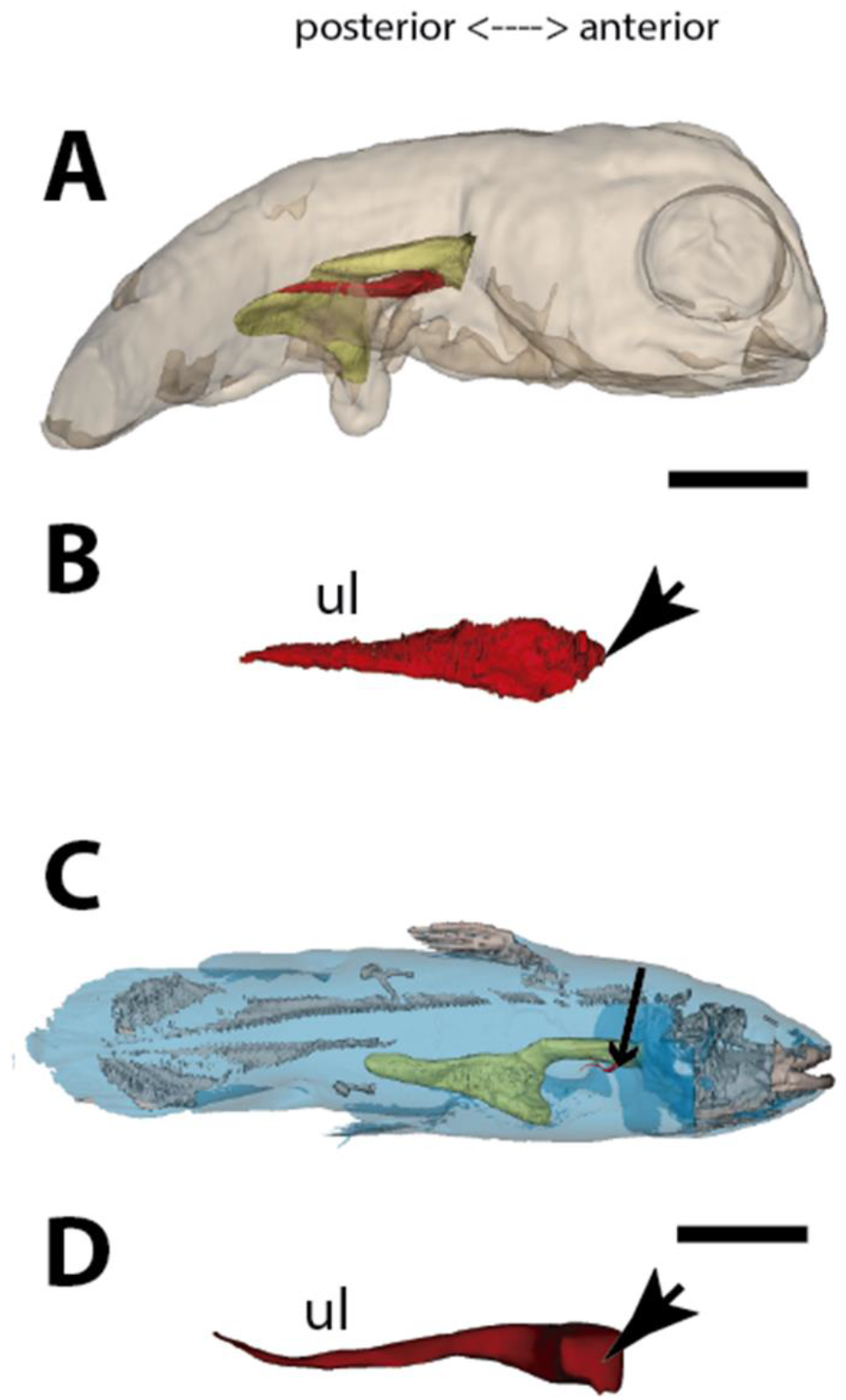
Three-dimensional reconstructions of the pulmonary complex of *Latimeria chalumnae*. **(A)** Early embryo of *Latimeria chalumane* (45 mm TL) in right lateral view (Cupello et al., 2015), **(B)** Isolated unpaired lung of the early embryo in dorsal view, **(C)** Adult specimen of *Latimeria chalumnae* (1300 mm TL) in right lateral view (Cupello et al., 2015), **(D)** Isolated unpaired lung of the adult specimen in dorsal view. Yellow, foregut including the stomach; red, lung. Arrowheads in **(B)** and **(D)** pointing to the lung connection to the foregut. Black arrow in **(C)** pointing to the lung. Ul, unpaired lung bud in **(B)** and unpaired lung in **(D)**. Scale bars, 5.0 mm **(A)**; 5.0 mm **(B)**; 200.0 mm **(C)**; 40 mm **(D)**. Modified from Cupello et al., 2015.

**Fig. 4.**
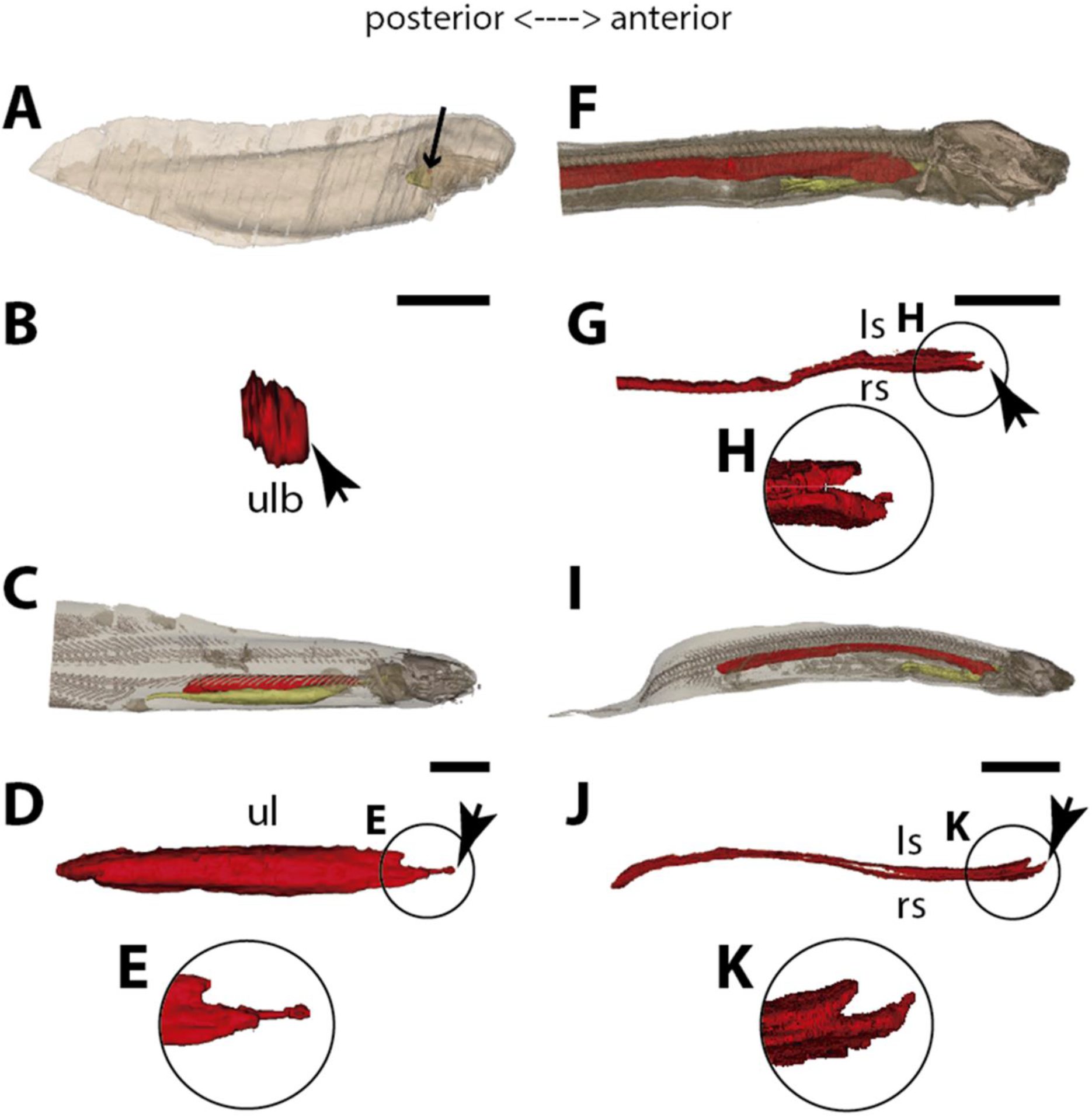
Three-dimensional reconstructions of the pulmonary complex of two species of Lepidosirenoidea. **(A)** Early embryo of *Neoceratodus forsteri* (13.5 mm TL) in right lateral view, **(B)** Isolated unpaired lung of the early embryo in dorsal view, **(C)** Adult specimen of *Neoceratodus forsteri* (200 mm TL) in right lateral view, **(D)** Isolated unpaired lung of the adult specimen in dorsal view, **(E)** Close-up of the lung unpaired connection to the foregut in (D), **(F)** Larva of *Lepidosiren paradoxa* (46 mm TL) in lateral view, **(G)** Isolated lung of the larval specimen in dorsal view, **(H)** Close-up of the lung unpaired connection to the foregut in (H), **(I)** Juvenile of *Lepidosiren paradoxa* young adult (68 mm TL) in lateral view, **(J)** Isolated lung of the juvenile specimen in dorsal view, **(K)** Close-up of the lung unpaired connection to the foregut in (J). Yellow, foregut including the stomach; red, lung. Black arrow in **(A)** pointing to the lung. Arrowheads in **(B)**, pointing to the lung connection to the foregut and in **(D)**, **(G)** and **(J)** pointing the pneumatic duct connection to the foregut. Ls, left sac; rs, right sac; ul, unpaired lung; ulb, unpaired lung bud. Scale bars, 2.5 mm **(A)**; 0.1 mm **(B)**; 20 mm **(C)**; 10 mm **(D, I, J);**5.0 mm **(F, G)**.

**Fig. 5.**
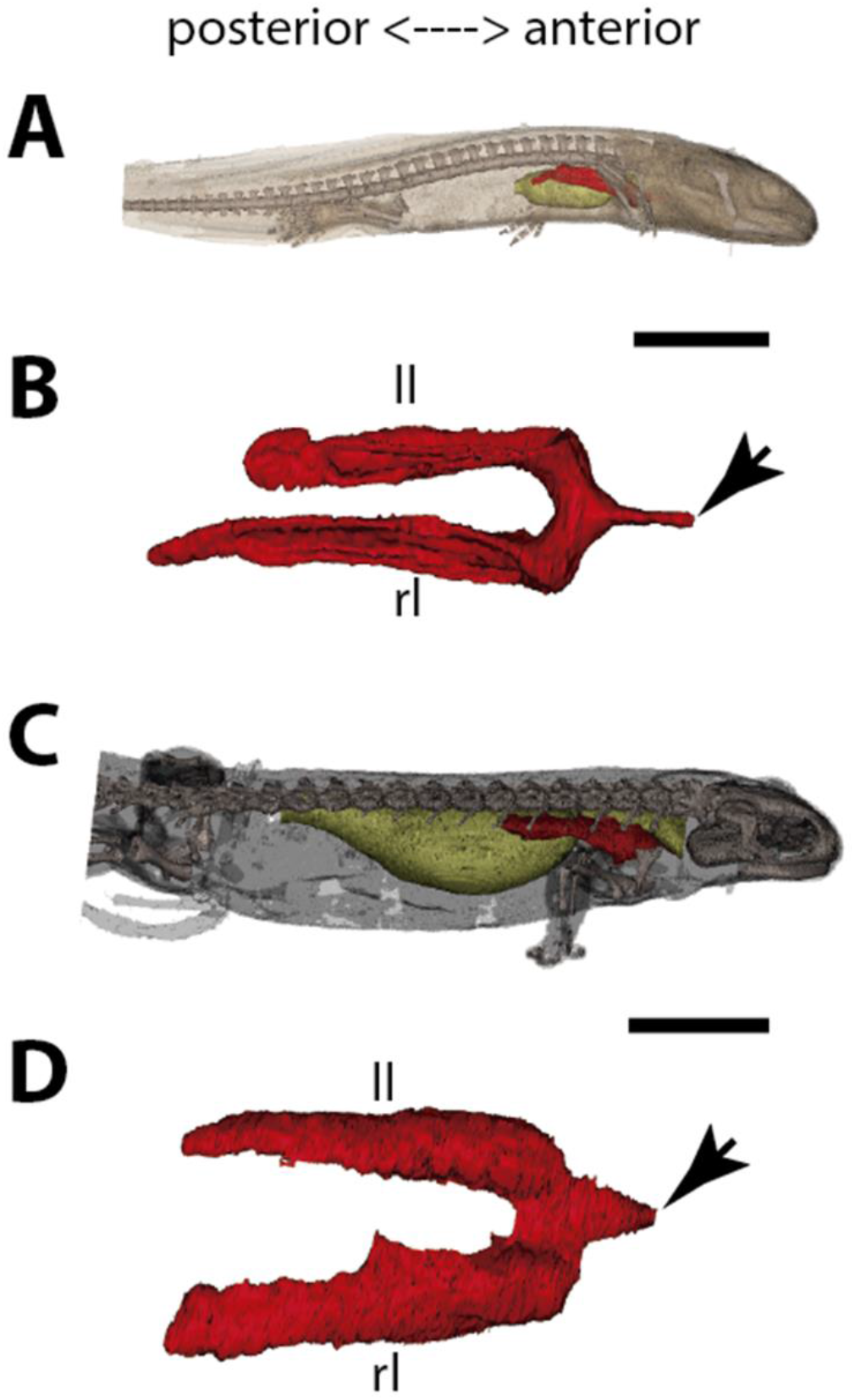
Three-dimensional reconstructions of the pulmonary complex of *Salamandra salamandra*. **(A)** Early larva of *Salamandra salamandra* (35.5 mm TL) in right lateral view, **(B)** Isolated paired lung of the larva embryo in dorsal view, **(C)** Juvenile of *Salamandra salamandra* (81.85 mm TL) in right lateral view, **(D)** Isolated unpaired lung of the juvenile specimen in dorsal view. Yellow, foregut including the stomach; red, lung. Arrowheads in **(B)** and **(D)** pointing to the trachea connection to the foregut. Ll, left lung; rl, right lung. Scale bars, 5.0 mm **(A)**; 3.125 mm **(B)**; 10 mm **(C)**; 6.25 cm **(D)**.

**Fig. 6.**
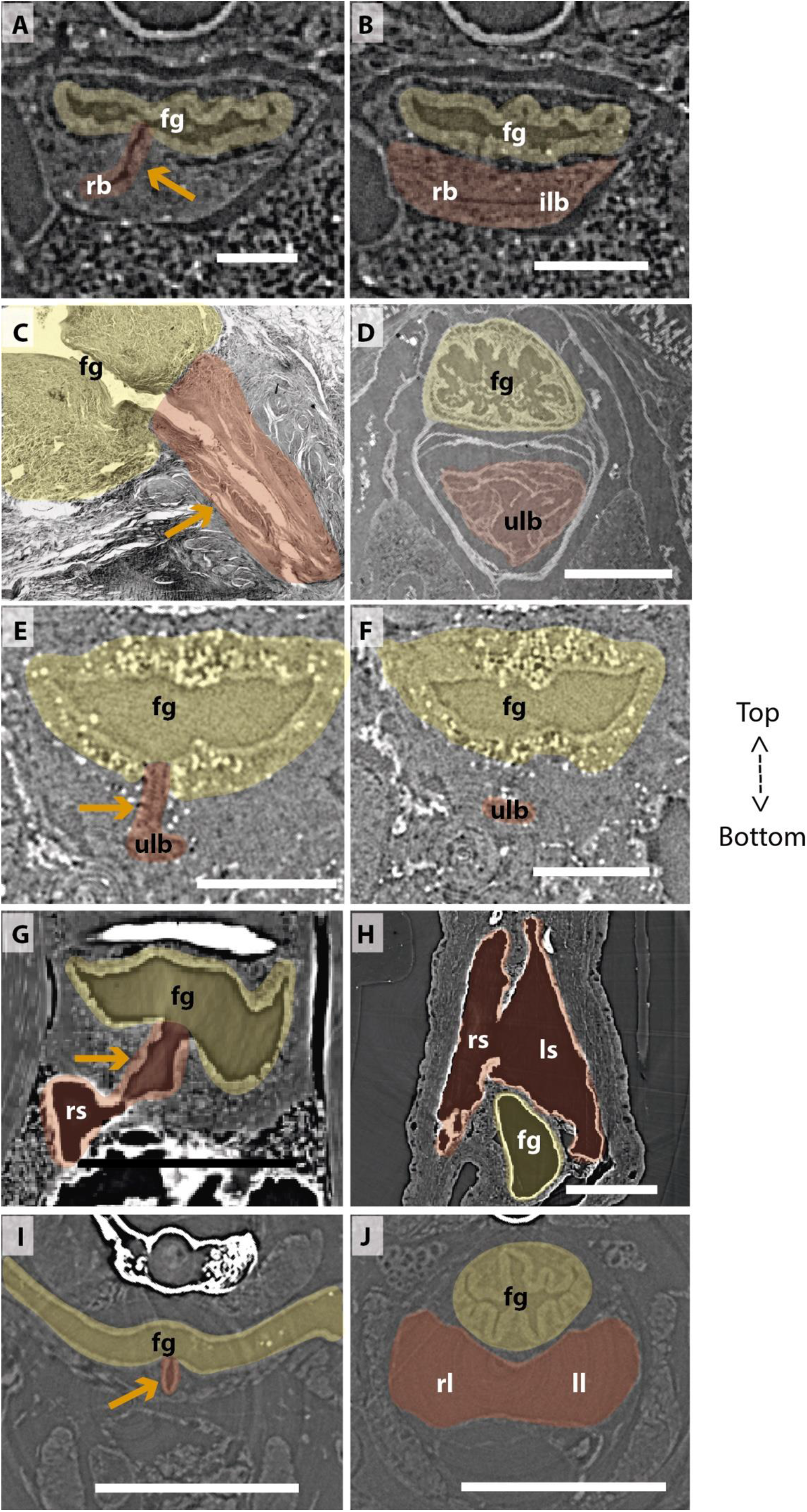
Comparison of sections showing the difference in lung origin and connection between unpaired (A-H) and true paired lungs (I, J). **(A, B)** Virtual section of *Polypterus senegalus* (12 mm TL), **(C)** Histological thin section of *Latimeria chalumnae* (127 cm) (Cupello et al., 2017a), **(D)** Virtual section of *Latimeria chalumnae* (40 mm TL) (modified from Cupello et al., 2017a), **(E, F)** Virtual section of *Neoceratodus forsteri* (16 mm TL), **(G, H)** Virtual section of *Lepidosiren paradoxa* (46 mm TL), **(I, J)** Virtual section of *Salamandra salamandra* (35.5 mm TL). Yellow, foregut including the stomach; red, lung. Orange arrows, opened connection between the foregut and the lung. Fg, foregut; ll, left lung; ls, left sac; rb, right bud; ilb, independent lung bud; rl, right lung; rs, right sac; ulb, unpaired lung bud. Scale bars, 0.25 mm **(A, B)**; 3.0 mm **(C)**; 1.0 mm **(D)**; 0.1 mm **(E, F)**; 0.5 mm **(G, H)**; 1.25 mm **(I, J)**.

## Results

### The lung development in *Polypterus senegalus*

From our observations on embryos of 8.0, 8.5, 9.1 and 9.3 mm TL (total length), the lung anlage develops as a ventral unpaired and tubular depression of the respiratory pharynx (the posterior portion of the pharynx), surrounded by undifferentiated mesenchymal cells^6^ (Fig. 1A, B, Fig. 2B, C). Only at the 12 mm TL larva, the left bud arises from the principal and primary lung anlage as a branch (Fig. 2D). Subsequently, the lung assumes its asymmetrical morphology, the left sac is smaller and remains ventral in the abdominal cavity, while the right one (principal tube) starts a partial turn up and stays parallel to the dorsal portion of the foregut (including the stomach). The left sac keeps a secondary connection to a lateral opening of the principal tube, and not to the foregut (Fig. 1 D, E, Fig. 6 A, B, Fig. S1, Fig. S2). Uundifferentiated dense cells surrounding the glottis are visible for the first time in specimens of 15.5 mm TL (black arrows in Fig. 2E). Airbreathing behavior starts at the juvenile stage in *P. senegalus* (2), and from juveniles of 23 mm TL onward, the blastema starts to develop into the muscular sphincter and respiratory epithelium at the glottis (ciliated cells intercalated by goblet cells). Right and left sacs are well developed and have a projection anterior to the connection with the small pneumatic duct in juveniles (Fig. 1D, E). The right tube is three times longer than the left one, with an expanded diameter in its caudal portion, posterior to the stomach (Fig. 1C, D).

The right and left sacs make a partial turn-up, remaining parallel to the dorsal surface of the upper gastrointestinal tract (one of each side). *Polypterus senegalus* lung is internally smooth and lacks alveolation at all the examined developmental stages, except for in the 45 mm TL juvenile, in which the most anterior projection of the lung, anterior to the connection with the pneumatic duct, is slightly compartmentalized. This evidence based on the first developmental stages of *P. senegalus* (embryos with 8.5 mm, 9.1 mm, 9.3 mm) lung prove that the lung bud initially develops as an unpaired anlage in this taxon (Fig. 1B, Fig. 2 B, C). The left sac develops secondarily from the right sac at later developmental stages, as a diverticulum, or a lobe, of the right primary lung (Fig. 2 E, F, Fig. S1, Fig. S2).

### The lung development in *Latimeria chalumnae*

Embryos of *L. chalumnae* display ventral compartmentalized unpaired lung throughout its length, suggesting alveolation (7), and in the early embryo (40 mm TL) a lateral and internal chamber is also present (Fig. 3; Fig. 6 D). At the latest developmental stages the pulmonary complex shows vestigial features, and no internal compartmentalization is recognizable (Lambertz and Perry, 2015). Adult specimens have constrictions and septations that divide the unpaired lung into separate lobes throughout its length, as in some fossil coelacanths (Cupello, Clément and Brito, 2019). Fossil coelacanths, from late Devonian to late Cretaceous, were most probably facultative air-breathers and made gas exchanges through their unpaired lungs and gills (Cupello, Clément and Brito, 2019; Brito et al., 2010). Although some authors suggest that *L. chalumnae* fatty organ evidences a paired lung, previous studies proved that this organ is not the second lung, since there is no opened connection between this organ and the foregut or lung, nor lung plates surrounding it (Cupello et al., 2015; Cupello et al., 2017a; Cupello et al., 2017b). Based on these, the paired condition of coelacanth lungs can be excluded.

### The lung development in *Neoceratodus forsteri*

The first developmental stage with lung anlage registered in this study is an early larva of 13.5 mm TL, with an unpaired morphology represented primarily by lung anterior projection (Fig. 4 B). In larvae of 16 mm, 17 mm, and 17.5 mm TL, although organogenesis is still not complete, a long and unpaired lung anlage is clearly identifiable and arises as a ventral depression of the post-pharyngeal foregut (Fig. 6 E, F). In the 19 mm TL specimen, the unpaired lung starts its dorsal turn up in relation to the dorsal portion of the foregut (including the stomach) (Fig. S3). From 20.5 mm TL onward, organogenesis is completed. In the larva of 20.5 mm TL, the lung remains unpaired and is ventrally connected to the post-pharyngeal foregut by a ventral, opened and long pneumatic duct. This organ has a projection anterior to the connection of the pneumatic duct and does not display alveolation/compartmentalization yet. According to previous studies (Kemp, 1982, 1986), air-breathing begins in *N. forsteri* at 25mm TL larval stage. Our results reveal that the larva with 26.5 mm TL presents a lung wall slightly pleated. From 50 mm TL larval stage onward, the lung wall is pleated eventually providing a high degree of alveolation. In the adult individual with 200 mm TL, the single lung displays internally two lateral chambers that are connected to a principal median chamber at both sides as a single structure (Fig. 4 C–E). In adult specimens of *N. forsteri,* the lung is highly compartmentalized by septa of smooth muscle and non-elastic connective tissue, as well as spongy alveolar structures (Grigg, 1965). At this developmental stage, the lung makes a complete dorsal turn-up at its posterior portion in relation to the gastrointestinal tract (Fig. 4 C). Although some authors pointed the presence of a second bud at the early developmental stages of *N. forsteri,* referred as the left lung (Spencer, 1893; Neumayer, 1904), the results presented herein show an indubitably unpaired configuration for *Neoceratodus* lung throughout the ontogeny (Fig. 4 A–E, Fig. 6 E, F, Fig. S3).

### The lung development in *Lepidosiren paradoxa*

Lungs of the four specimens studied herein, from larva to adults (larva with 46 mm TL, juveniles with 68 mm TL and 222.1 mm TL, and adult with 400 mm TL), display a similar morphology and, surprisingly, left and right tubes do not arise simultaneously. Only the right sac is connected to the pharynx by a long pneumatic duct (Fig. 6 G, H, Fig. S4). The left sac is a branch of the right one, connected by a posterior and secondary opening at the lung level, already in dorsal position in relation to the foregut (Fig. 6 G, H, Fig. S4). There is no connection of the left sac with the pneumatic duct. In *L. paradoxa,* only the pneumatic duct is ventrally positioned, and the lung makes a complete dorsal turn up from the right side of the upper gastrointestinal tract (Fig. 4 F–K, Fig. S5), just after the ventral connection to the pharynx. This complete dorsal turn-up is also seen in the lung of adult specimens of *N. forsteri* (Fig. 4 I–K). There are no anterior projections of the lung. Lung compartmentalization is clearly observable through dissections, evidencing the high degree of alveolation (Fig. S5). Our results indicate that the lung of *L. paradoxa* is, in fact, remarkably similar to *P. senegalus* lung. The so-called left lung of *L. paradoxa* is most likely a diverticulum or a modified lateral lobe, which had evolved secondarily, an advantage for enlarging the surface area for oxygen-uptake, eventually enabling the obligatory air-breathing performance in the linage towards *L. paradoxa*.

### The lung development in *Salamandra salamandra*

In early larvae with 35.5 mm TL and 42.8 mm TL, paired lungs are collapsed in its middle and posterior portion (Fig. 5 A, B). The internal lung wall is thin and smooth, without compartmentalization and/or alveolation in its inner wall (Fig. 6 J, Fig. S6). From the early larvae onward, the muscular glottis develops on the ventral portion of the pharynx, and both left and right lungs arise simultaneously and symmetrically from a long trachea and paired first order bronchioles (Fig. 5 C, D, Fig. 6 I, J, Fig. S6). Lungs are symmetrical in size and morphology and are placed in the anteriormost portion of the abdominal cavity, as described for other tetrapods (Fig. 5 C, D).

In postmetamorphic juveniles (of 81.85 mm TL), paired lungs are already functional, not collapsed, and the main organ for oxygen-uptake (Goniakowska-Witalińska, 1978, 1982).

From this developmental stage onward, lungs are highly compartmentalized by multiple septa. Due to the paired and compartmentalized anatomy, the lung surface area for oxygen-uptake, as well as its volume capacity, increase substantially – both important features for a functional lung in dry environments. Here we confirm that at different developmental stages of *S. salamandra*, lungs are truly paired since both left and right lungs arise simultaneously and symmetrically and are directly connected to the trachea. Throughout the ontogeny, *S. salamandra* lungs have a ventral origin, and makes a partial dorsal turn-up in its posterior portion, remaining parallel to the dorsal wall of the upper gastrointestinal tract. Due to the rarity of soft tissue preservation in the fossil record, only one species of salamandrid, *Phosphotriton sigei* presents its lung preserved (Tissier, Rage and Laurin, 2017). This lung is described as multichambered, placed in the anteriormost part of the abdominal cavity (Tissier, Rage and Laurin, 2017), such as in the living salamandrid described above.

## Discussion

Traditionally, vertebrate lungs are defined as ventral paired organs derived from the ventral portion of the posterior pharynx or post-pharyngeal foregut (Perry et al., 2001; Funk, Lencer and McCune, 2020; Lambertz and Perry, 2015; Graham, 1997; Kardong, 2015). However, we demonstrate here the presence of an unambiguous unpaired lung, that develop from the ventral foregut, but sometimes occupy the dorsal position later in the development of osteichthyan fishes (Fig. 7). Based on extensive developmental series of different vertebrate taxa, we present a new interpretation of some lungs previously considered as paired and, therefore, a new definition of paired lungs. Based on our results, true paired lungs are stated when bilateral lung buds arise simultaneously and are both connected directly to the foregut, as observed in the salamander (Fig. 7).

**Fig. 7.**
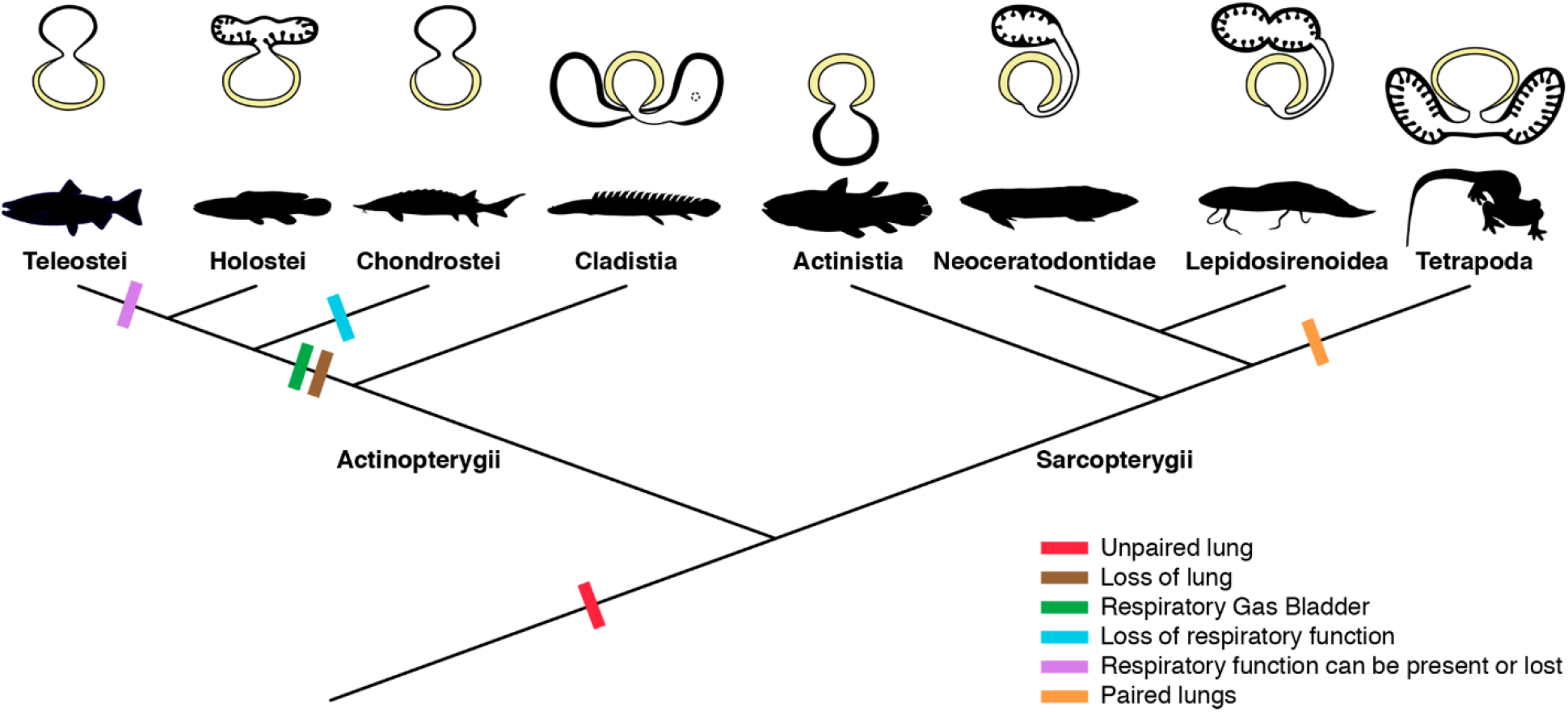
Schematic figure reconstructing the evolutionary history of vertebrate lungs. All living actinopterygian and sarcopterygian fishes have unpaired lungs. True paired lungs are a synapomorphy of tetrapods. Dashed circle in Cladistia lung pointing to the secondary and independent opening to a left sac, at the lung level. Modified from Liem, 1988. This figure was made with free silhouettes from PhyloPic.

The sister group to all other living actinopterygian (polypterids) and all living sarcopterygian fishes have a clear unpaired lung in early developmental stages that can be developed in later stages either in a unilobed or a secondarily multilobed lung (Fig. 7), and not in a true paired lung. *Polypterus senegalus* and the lungfish *L. paradoxa* possess secondary multilobed structure from the larval stage onward since the lung is derived from a unilateral connection to the foregut. The presence of this secondary multilobed morphology is an advantage for the obligatory air-breathing behavior of these taxa, raising the respiratory compliance. In the teleost *Batrachomoeus trispinosus*, the non-respiratory gas bladder is also described as paired (Rice and Bass, 2009), although it is certainly a secondary condition.

The most parsimonious scenario inferred from our data mapped on the phylogenetic framework (Fig. 7) is that the vertebrate lung was unpaired at the evolutionary origin. Since soft tissue are rarely preserved in fossils, living lunged osteichthyans are key taxa for the understanding of how evolutionary constraints shaped breathing adaptations on land. Our study revealed that the ancestral condition of the lung is a median unpaired organ (Fig. 7), thereby being inconsistent with the scenario that the lung evolved through a modification of the posteriormost pharyngeal pouch assumed to be present in primitive taxa (Kastschenko, 1887; Goodrich, 1931). Consequently, the evolutionary origin of the lung was likely independent of the pharyngeal pouch at the morphological level.

From this evolutionary point of view, we complement lung definition as an unpaired or paired respiratory organ, or its vestigial form, that develops and are ventrally connected to the foregut. Some criteria previously used for discriminating lungs from gas bladders are no longer useful, including paired/unpaired organization, position ventral to the alimentary tract (Marcus, 1937; Funk, Lencer and McCune, 2020; Lambertz and Perry, 2015; Graham, 1997), as well as its function. The dorsal position of the majority of osteichthyans lungs described here may be related to its dual and secondary functionality of respiration and buoyancy control (Thomson, 1968). Actually, the only morphological characteristic that can be used to distinguish lungs and gas bladders is the ventral and dorsal origins from the foregut, respectively (Funk, Lencer and McCune, 2020; Cass, Servetnick and McCune, 2013). This phenotypic differentiation into true paired lungs in tetrapods may be related to differential gene expressions (Funk, Lencer and McCune, 2020; Bi et al., 2021). Nevertheless, at the developmental genetic level, the possibility of co-options of gene regulatory networks of the pharyngeal pouch morphogenesis cannot be excluded, as both the lung bud and pharyngeal pouch develop through the invagination of the foregut endoderm. Our results open the door for future molecular analyses to trace possible regulatory elements for the evolutionary transition from unpaired lungs to true paired lungs in tetrapods.

According to morphological evidence presented here, bifurcation morphogenesis into true paired lungs was not developed yet in osteichthyan fish ancestors. The bilaterally paired nature of the lung evolved only in the lineage towards the tetrapods, as a synapomorphy of this clade (Fig. 7). This morphological modification brought about improvement of the efficiency in oxygen-uptake from the air, as the paired lungs having parallel air flows exchange the air more quickly than the unpaired lung having only single air flow does. This innovation led to the elevation of metabolic rate that was required for the sustained body support against the gravity. Paired lungs may have been present also in early tetrapods and were probably essential to raise lung surface area and volume capacity during the evolution of vertebrate respiratory system and the air-breathing intensification at the water to land transition.

## Materials and Methods

### Specimens information

All specimens used in this work are permanently housed in collections of public institutions. No specimens were collected alive in the field for this work. *Polypterus senegalus* specimens were originally obtained for the study on the molecular developmental in polypterids (Tatsumi et al., 2016). Nine specimens here studied are: six late embryos (free embryonic phase or postembryos) of 8.0 mm TL (PS-001-01) and histological thin-section of another specimen of 8.0 mm TL (PSS-No1), 8.5 mm TL (PS-001-02), 9.1 mm TL (PSS-No2) and 9.3 mm TL (two specimens, PS-001-03); four larva of 12 mm TL (two specimens, PS-001-04), 15.5 mm TL (PSS-No3) and 18.0 mm TL (PSS-No4); and three juveniles of 20 mm TL (PS-001-05), 23 mm TL (PS-001-06), and 45 mm TL (PS-001-07). We indicate the developmental stages (embryo, larvae, juveniles, and adults) following Bartsch, Gemballa and Piotrowski (1997). Specimens and histological material are housed at the Department of Anatomy of the Jikei University School of Medicine (Tokyo, Japan).

Four specimens of *Lepidosiren paradoxa* here studied are from the collections of the Universidade do Estado do Rio de Janeiro and were collected legally in 2008, with the permission number 11471-1. The specimens are registered under the acronym UERJ-PN: UERJ-PN 550 is a larva of 46 mm TL; UERJ-PN 262 is a juvenile of 68 mm TL; UERJ-PN 238 is juvenile of 222.1 mm TL; and PC02 is an adult of 400 mm TL. We follow Kerr (1900) for the developmental staging of *Lepidosiren*.

Specimens of *Neoceratodus forsteri* were collected legally from Department of Biological Sciences, Macquarie University, Sydney, Australia, and transported with the permission of CITES (Certificate No. 2009-AU-564836). The developmental series comprises fourteen specimens. Sizes are: an early embryo of 13.5 mm TL (IMU-RU-SI-0013); 11 larvae of 16 mm TL (IMU-RU-SI-0017), 17 mm TL (IMU-RU-SI-0019 and IMU-RU-SI-0022], 17.5 mm TL (IMU-RU-SI-0037), 19 mm TL (IMU-RU-SI-0038), 20.5 mm TL (IMU-RU-SI-0039), 24 mm TL (IMU-RU-SI-0040), 25.5 mm TL (IMU-RU-SI-0041), 26.5 mm TL (IMU-RU-SI-0042), 30 mm TL (IMU-RU-SI-0043), and 50 mm TL (IMU-RU-SI-0045); a juvenile of 70 mm TL (IMU-RU-SI-0048); and an adult specimen of 200 mm TL (KPM-NI 11384). For the developmental identification (embryos, hatchlings/larvae, juveniles, and adults) we follow Kemp (1982, 2011) and Ziermann et al. (2018).

Six *Salamandra salamandra* specimens were obtained on loan at the amphibian collection of the Muséum national d’Histoire naturelle (Paris, France). The three developmental stages (as described at the MNHN collection) are: two early larvae MNHN 1978.636 (1) of 35.5 mm TL and MNHN 1978.636 (2) of 42.8 mm TL; larva MNHN 1985.9039 of 49.6 mm; larva in metamorphosis MNHN 1978.542 of 54.44 mm TL; small juvenile MNHN 1988.7177 of 50 mm TL; juvenile MNHN 1962.1004 of 81.85 mm TL.

Institutional abbreviations: IMU-RU-SI, Iwate Medical University, Ryozi Ura Collection, Japan; PS, *Polypterus senegalus;* PSS, *Polypterus senegalus* sections; KPM-NI, Kanagawa Prefectural Museum Natural History, Odawara, Japan; MNHN, Muséum national d’Histoire naturelle, Paris, France; UERJ-PN, Universidade do Estado do Rio de Janeiro, Peixes Neotropicais.

### X-ray tomography

Due to the extremely small size of the embryos and larvae, and to the weak density difference between soft tissues of the abdominal cavity, propagation phase-contrast microtomography was the unique way to study their anatomy and histology at micrometer scale. Phase-contrast microtomography being only achieved at synchrotron sources, we accessed the anatomy of these rare and tiny samples at the Synchrotron SOLEIL and Synchrotron SPring-8. The high brightness of the synchrotrons was essential for our material and enabled the collection high resolution scans in short timescales.

Specimens of *P. senegalus*, *L. paradoxa* and *S. salamandra* were imaged at the PSICHÉ beamline of the SOLEIL Synchrotron (Saint-Aubin, France) while *N. forsteri* specimens were scanned at SPring-8 Synchrotron. The specimens were scanned isolated in a plastic tube filled with Phosphate-buffered saline (PBS) for *P. senegalus* and *N. forsteri*, ethanol for *L. paradoxa* and formaldehyde for *S. salamandra*. They were immobilized in vertical position using gauze pads, and/or sank inside the tip of a plastic pipette in the case of tiny individuals, in order to benefit as much possible from the available field of view and thus achieve the highest possible resolution.

At SOLEIL Synchrotron, imaging was performed using a monochromatic beam with an energy of 25 keV. A series of acquisitions with vertical movement of the sample were recorded to extend vertically the field of view and image the entire (or most of the) individual. Two distinct setups were used to accommodate the different sizes of the individuals (size variations occurring both between developmental stages and taxa). (1) Small individuals were scanned using a field of view of 2.6 × 2.6 mm^2^ (5x magnification) resulting in a projected pixel size of 1.3 μm, and a propagation distance of 148 mm. (2) Larger individuals were scanned using a field of view of ~12.6 × 3.3 mm^2^ (1x magnification) resulting in a projected pixel size of 6.17 μm, and a propagation distance of 500 mm. For individuals slightly wider than these field of views, the latter were extended horizontally by positioning the rotation axis off-centre and acquiring data over a 360° rotation of the sample. The volumes were reconstructed from the (vertically) combined radiographs using PyHST2 software (Mirone et al., 2014), with a Paganin phase retrieval algorithm (Paganin et al., 2002). The huge resulting volumes (from 70 Gb to 1.2 Tb) were reduced (crop, rescale 8-bit, binning) to facilitate 3D data processing.

Specimens of *Neoceratodus forsteri* (from 13.5 to 70 mm TL) were imaged at the SPring-8 Synchrotron, beamline 20B2. For specimens from 13.5 mm TL to 30 mm TL, a beam energy of 15 keV was used with a double bounce Si (111) monochromator. Data were obtained at three different resolutions, and correspondingly used three combinations of two lenses and fluorescent material, as follows, 2.75 μm/voxel; 1st-stage lens: “beam monitor 2” f35 mm; 2nd-stage lens: Nikon 85 mm lens; GADOX thickness: 15 μm 4.47 μm/voxel; 1st-stage lens: “beam monitor 2” f35 mm; 2nd-stage lens: Nikon 50 mm lens; GADOX thickness: 15 μm 12.56 μm/voxel; 1st-stage lens: “beam monitor 5” f200 mm; 2nd-stage lens: Nikon 105 mm lens; GADOX thickness: 25 μm.

Datasets were acquired at propagation distances of 2.75 μm/voxel, 4.47 μm/voxel: 600 mm; 12.56 μm/voxel: 3 m and three different exposure times of 70 ms, 150 ms, and 200 ms per projection. Field of view were: pixel size × 2048 (2.75 × 2048 = 5632 μm; 4.47 × 2048 = 9154.56 μm; 12.56 × 2048 = 25722.88 μm) A total of 1800 projections were recorded per scan as the sample was rotated through 180°. A high-resolution computerizedaxial tomography scanning (CAT scan) was performed for the adult specimen of *Neoceratodus* (KPM-NI 11384) of 200 mm TL at the National Museum of Nature and Science (Tokyo, Japan) using the following scanning parameters: effective energy 189 kV, current 200 mA, voxel size 9.765 μm and 1000 views (slice width 0.1 mm).

### Segmentation and three-dimensional rendering

Segmentation and 3D rendering were performed using the software MIMICS Innovation Suite 20.0 (Materialise) at the Laboratório de Ictiologia Tempo e Espaço of the Universidade do Estado do Rio de Janeiro.

## Acknowledgments

We thank J. Joss (Macquarie University) for providing *Neoceratodus* embryos. We are grateful to Dr H. Seno (curator of the Ichthyology Collection of the Kanagawa Prefectural Museum of Natural History) for the loan of *Neoceratodus* specimen KPM-NI 11384.

## Funding

Coordenação de Aperfeiçoamento de Pessoal de Nível Superior—Brasil (CAPES)—grant Finance Code 001 (Programa Nacional de Pós Doutorado-PNPD) (CC)

Programa de Apoio à Docência (PAPD) grant E-26/007/10661/2019)—Universidade do Estado do Rio de Janeiro (CC)

Interdisciplinary Collaborative Research Program of the Atmosphere and Ocean Research Institute, the University of Tokyo JP (CC, TS)

Prociência fellowship CNPq grant 310101/2017-4 (PMB)

FAPERJ grant E-26/ 202.890/2018 (PMB)

## Author contributions

Examples:

Conceptualization: CC, YY, PMB

Synchrotron acquisitions: CC, TH, YY, PG, SI, AK, MH, KU, PMB

CT scan acquisitions: RM

Computerized microtomography rendering: CC, NT

Histological thin sections preparation: NT, MO

Specimens dissection: CC, NT, SI

Tomographic setups and data processing: PG, RM, AK, MH, KU

Figures: CC, PG

Data interpretation and Writing—original draft: CC, TH, NT, YY, PG, MO, PMB

Writing—final writing & manuscript approval: CC, TH, NT, YY, PG, SI, RM, TS, AK, MH, KU, MO, PMB

## Competing interests

All authors declare no competing interests.

## Data and materials availability

All data are available in the main text or the supplementary materials.

## Supplementary Materials

Supplementary information is available for this paper at https://doi.org/.

## Supplementary Information

**Fig. S1.**
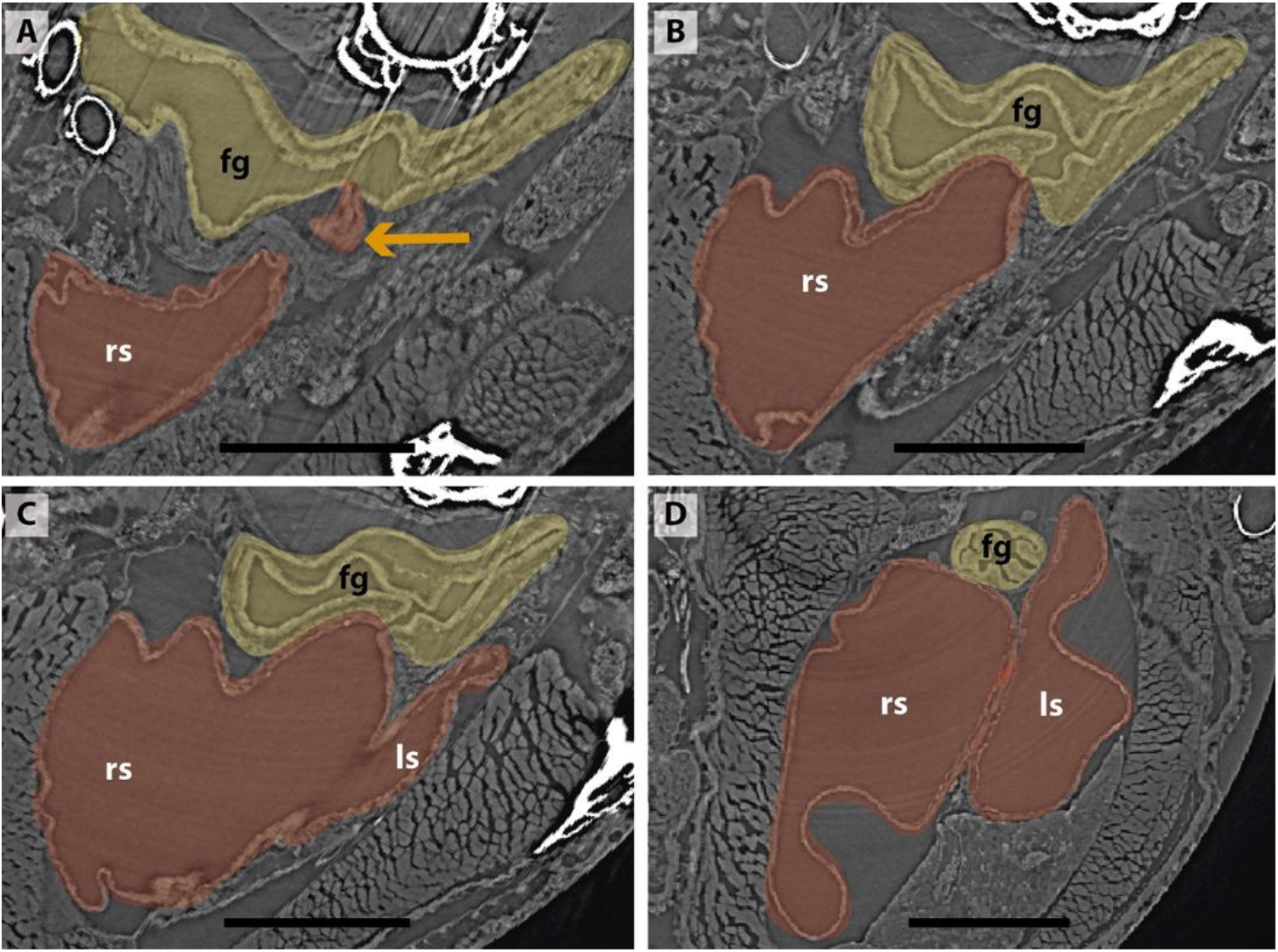
Sections of synchrotron X-ray microtomography of a juvenile of *Polypterus senegalus* (23 mm TL). **(A)** Unpaired lung origin. **(B)** Right sac arising from the foregut. **(C)** Left sac arising from an independent and lateral connection to the right sac. **(D)** Right and left sacs. Yellow, foregut; red, lung. Orange arrow, opened connection between foregut and lung. Fg, foregut; ls, left sac; rs, right sac. Scale bars, 0.5 mm **(A-D)**.

**Fig. S2.**
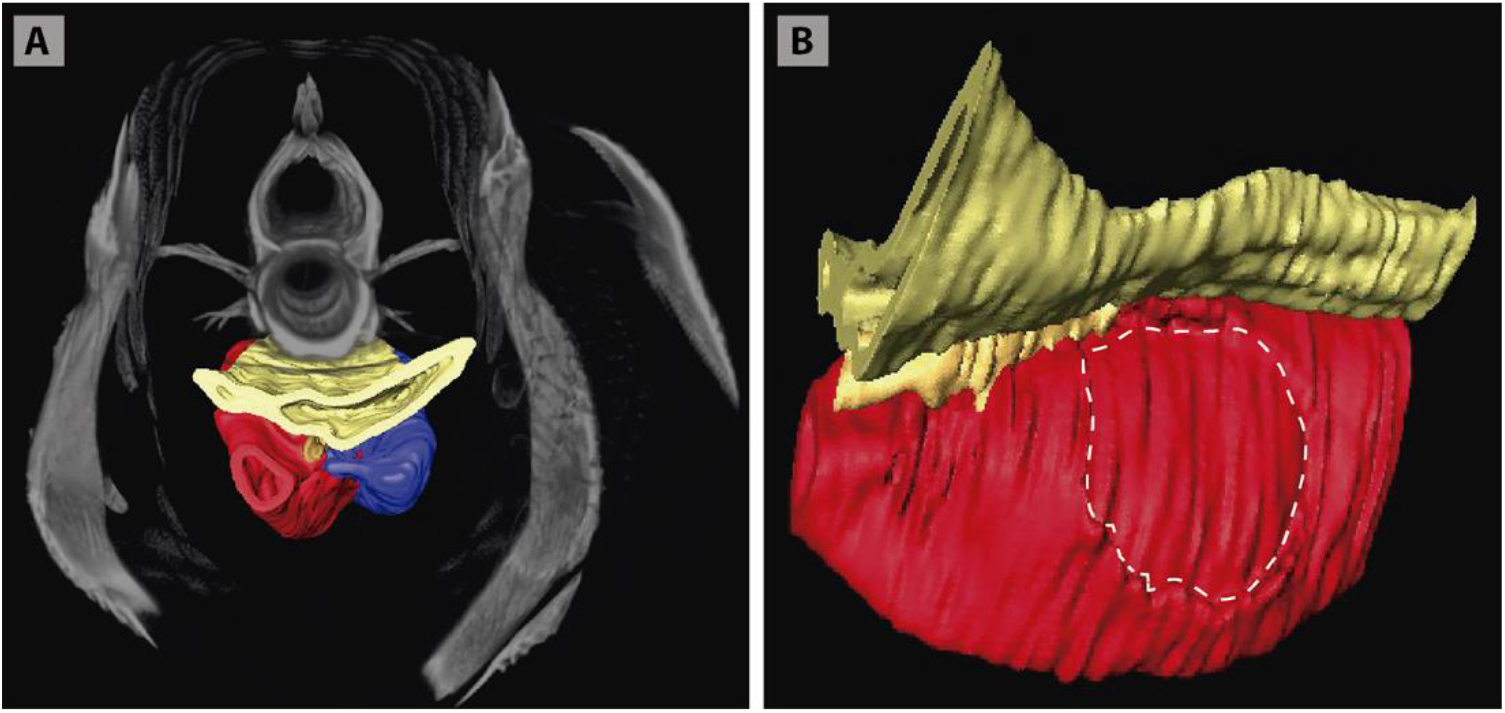
Three-dimensional reconstructions of the pulmonary complex of *Polypterus senegalus*. **(A)** Virtual section of the juvenile (45 mm TL) in anterior view, evidencing the oesophagus and the lung in 3D. **(B)** Isolated right lung of the juvenile in lateral view, evidencing the independent and secondary connection of the left sac to the right one by a lateral opening. Yellow, foregut including the stomach; red, right sac; blue, left sac, dashed line, independent and secondary connection of the left sac to the right one.

**Fig. S3.**
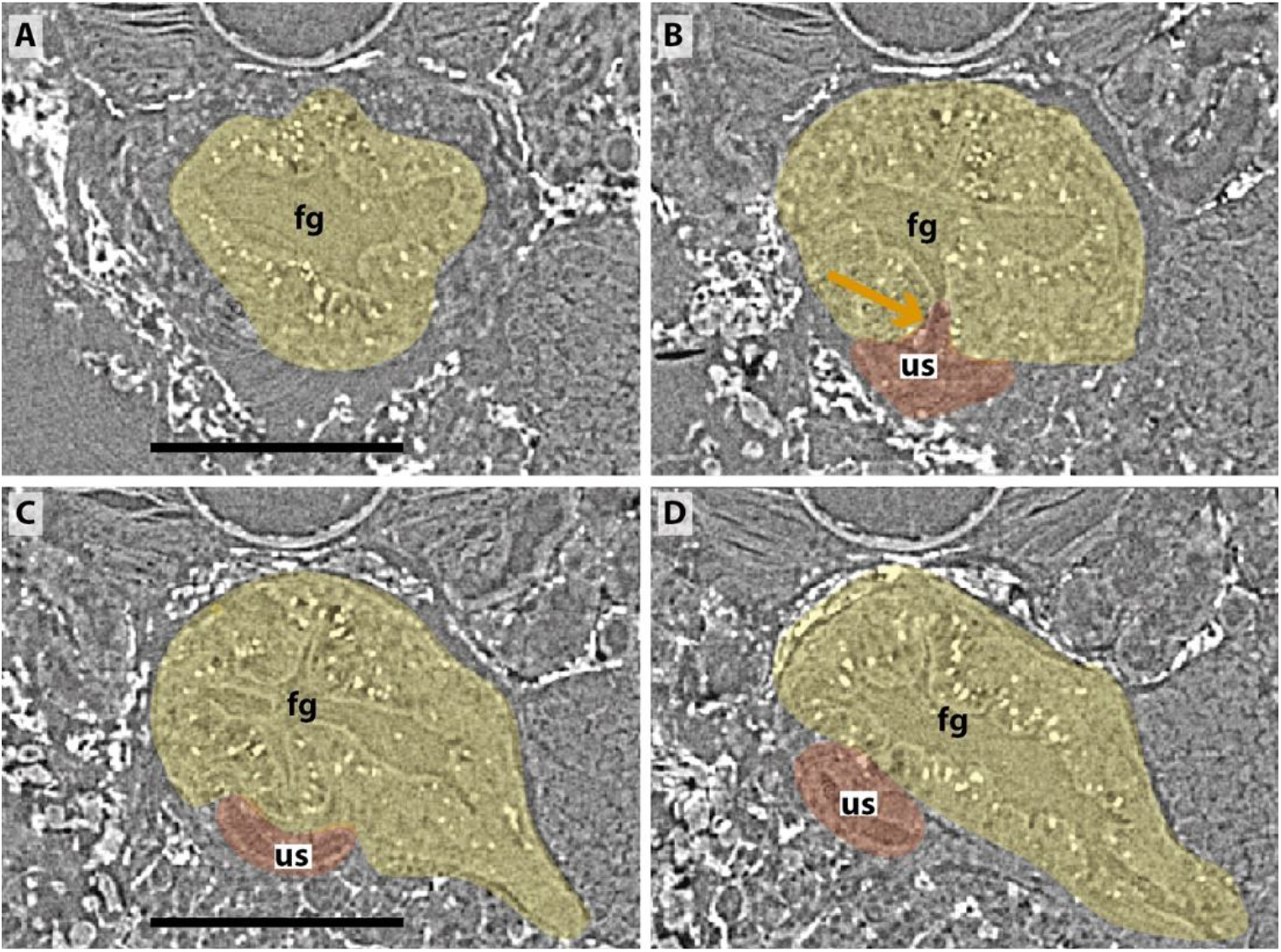
Sections of synchrotron X-ray microtomography of a larva of *Neoceratodus forsteri* (19 mm TL). **(A)** Unpaired lung origin. **(B)** Unique sac arising from the foregut. **(C, D)** Unique sac developing. Yellow, foregut; red, lung. Orange arrow, opened connection between foregut and lung. fg, foregut; us, unique sac. Scale bars, 0.5 mm **(A-D)**.

**Fig. S4.**
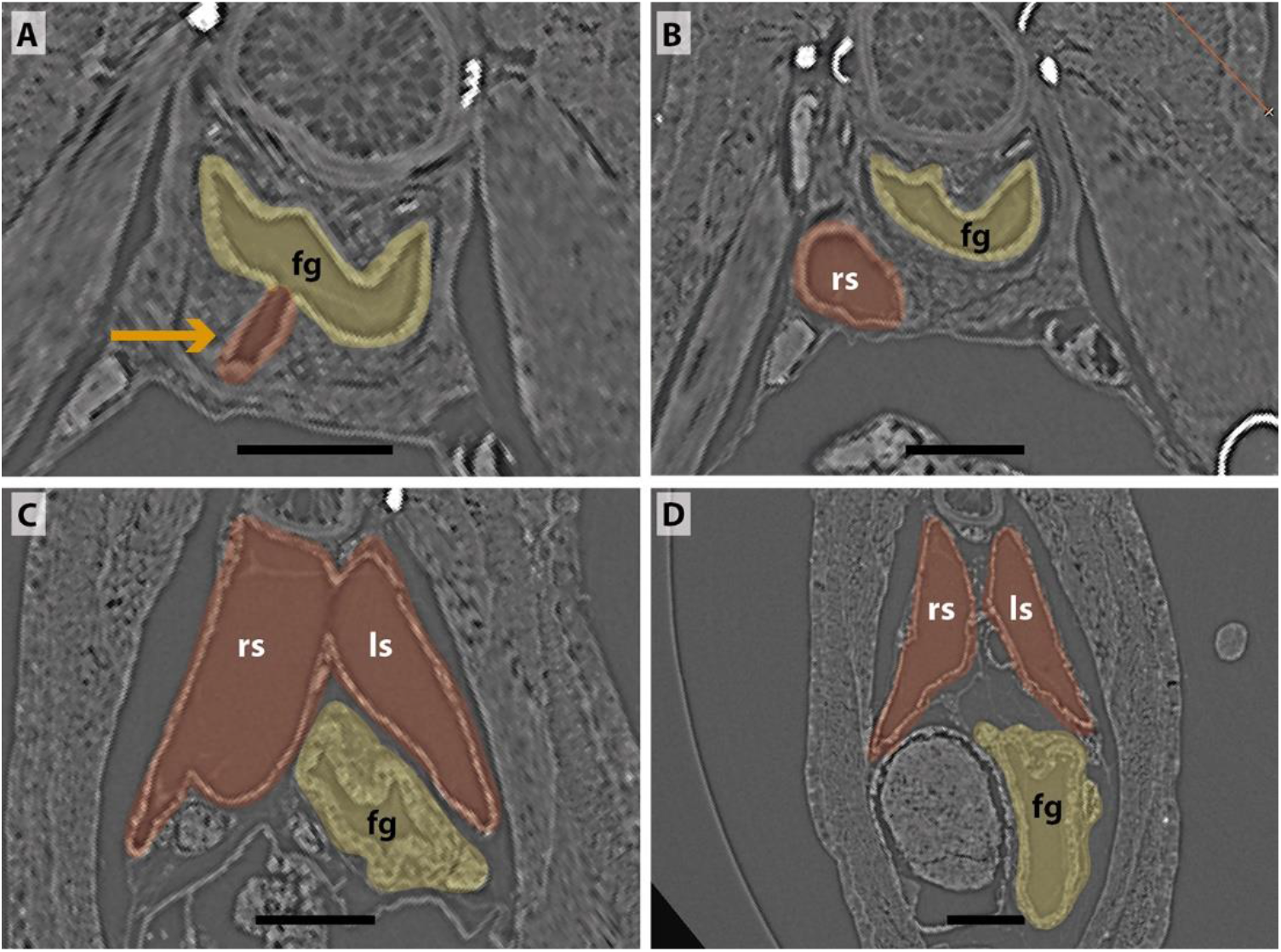
Sections of synchrotron X-ray microtomography of a juvenile of *Lepidosiren paradoxa* (68 mm TL). **(A)** Unpaired lung origin. **(B)** Right sac arising from the foregut. **(C)** Left sac arising from an independent and lateral connection to the right sac. **(D)** Right and left sacs. Yellow, foregut; red, lung. Orange arrow, opened connection between foregut and lung. fg, foregut; ls, left sac; rs, right sac. Scale bars, 0.5 mm **(A-D)**.

**Fig. S5.**
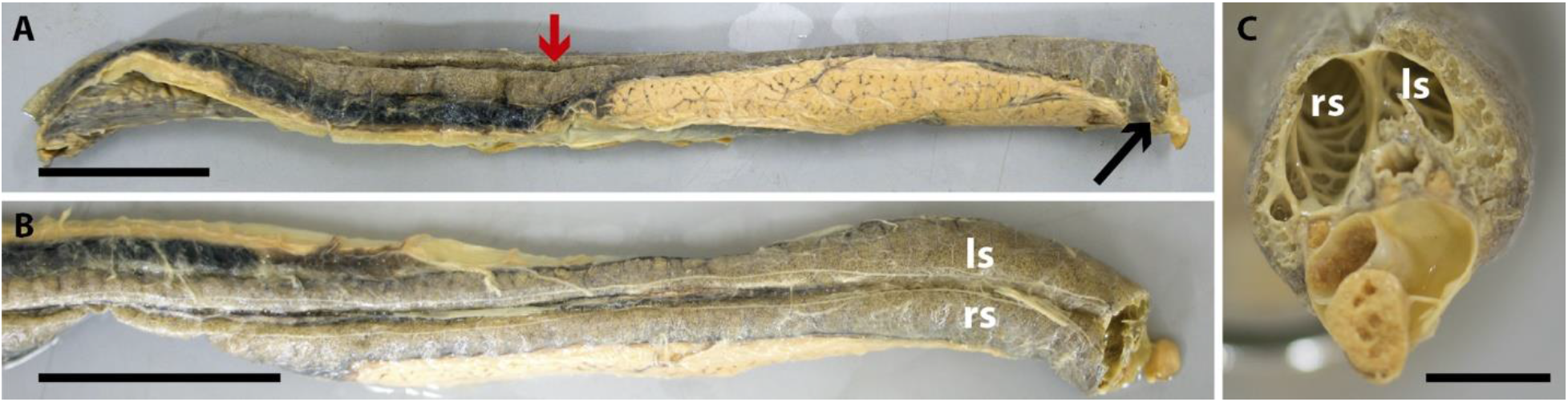
Dissection of the lung of an adult *Lepidosiren paradoxa* (400 mm TL). Red arrow, lung. Black arrow, ventral insertion of the right sac. ls, left sac; rs, right sac. Scale bars, 50 mm **(A, B)**; 10 mm **(C)**.

**Fig. S6.**
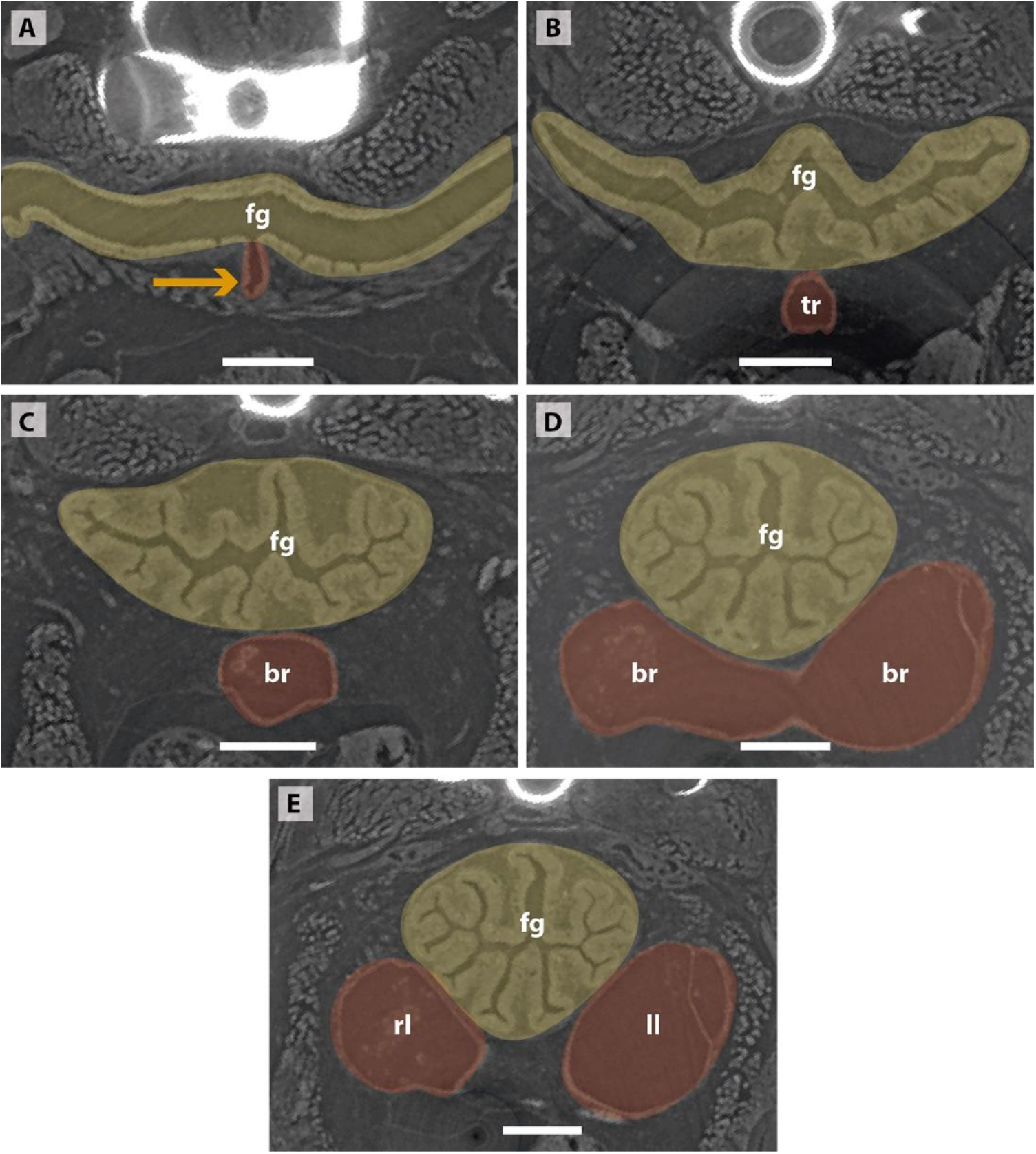
Sections of synchrotron X-ray microtomography of a larva of *Salamandra salamandra* (42.8 mm TL). **(A, B)** Trachea arising. **(C, D)** Fist order bronchioles. **(E)** Right and left lungs arising simultaneously and symmetrically. Yellow, foregut; red, lung. Orange arrow, opened connection from the foregut. br, braonchile; fg, foregut; ll, left lung; rl, right lung; tr, trachea. Scale bars, 0.5 mm **(A-D)**.

## Notes

### Competing Interest Statement

The authors have declared no competing interest.

